# Evidence accumulation and associated error-related brain activity as computationally-informed prospective predictors of substance use in emerging adulthood

**DOI:** 10.1101/2020.03.06.981035

**Authors:** Alexander S. Weigard, Sarah J. Brislin, Lora M. Cope, Jillian E. Hardee, Meghan E. Martz, Alexander Ly, Robert A. Zucker, Chandra Sripada, Mary M. Heitzeg

## Abstract

**Rationale:** Substance use peaks during the developmental period known as emerging adulthood (ages 18–25), but not every individual who uses substances during this period engages in frequent or problematic use. Although individual differences in neurocognition appear to predict use severity, mechanistic neurocognitive risk factors with clear links to both behavior and neural circuitry have yet to be identified. Here we aim to do so with an approach rooted in computational psychiatry, an emerging field in which formal models are used to identify candidate biobehavioral dimensions that confer risk for psychopathology.

**Objectives:** We test whether lower efficiency of evidence accumulation (EEA), a computationally-characterized individual difference variable that drives performance on the go/no-go and other neurocognitive tasks, is a risk factor for substance use in emerging adults.

**Methods and Results:** In an fMRI substudy within a sociobehavioral longitudinal study (*n*=106), we find that lower EEA and reductions in a robust neural-level correlate of EEA (error-related activations in salience network structures) measured at ages 18–21 are both prospectively related to greater substance use during ages 22–26, even after adjusting for other well-known risk factors. Results from Bayesian model comparisons corroborated inferences from conventional hypothesis testing and provided evidence that both EEA and its neuroimaging correlates contain unique predictive information about substance use involvement.

**Conclusions:** These findings highlight EEA as a computationally-characterized neurocognitive risk factor for substance use during a critical developmental period, with clear links to both neuroimaging measures and well-established formal theories of brain function.

## 1. Introduction

Problematic substance use, which affects tens of millions of Americans, has serious consequences and exacts a significant physical, emotional, and financial burden (Centers for Disease Control and Prevention, 2016, 2018; Florence et al., 2016; National Drug Intelligence Center, 2011; Xu et al., 2015). However, not everyone who tries alcohol, tobacco, or other drugs goes on to develop a substance use disorder (SUD). A better understanding of the factors— including behavioral, cognitive, and neural variables—that predict problematic substance use is therefore essential for improving prevention and treatment efforts.

Substance use typically increases throughout adolescence and peaks during individuals’ early-to-mid 20s, in a period characterized as “emerging adulthood” (Arnett, 2000; Schulenberg et al., 2019). As neural circuitry involved in the control of cognition and actions continues to mature throughout these developmental periods, immaturity in this circuitry is theorized to contribute to problematic substance use (Casey et al., 2011; Schulenberg et al., 2016; Shulman et al., 2016). Consistent with these assertions, behavioral and neural probes of “inhibitory control”, a neurocognitive function that is proposed to allow individuals to stop dominant or prepotent responses (Miyake et al., 2000), appear to relate to substance use behaviors. This construct is often measured using the go/no-go task (Figure 1a), in which participants are instructed to make a response following certain stimuli (“go” trials) but to inhibit their response following others (“no-go” trials). Meta-analytic studies have found poorer behavioral performance on the go/no-go across multiple substance use disorders (Smith et al., 2014). Crucially, longitudinal neuroimaging studies have found that reduced go/no-go task-related activation in prefrontal cortical regions prospectively predicts problem use in both early (Heitzeg et al., 2014) and later adolescence (Mahmood et al., 2013; Norman et al., 2011; Wetherill et al., 2013).

**Fig. 1.**
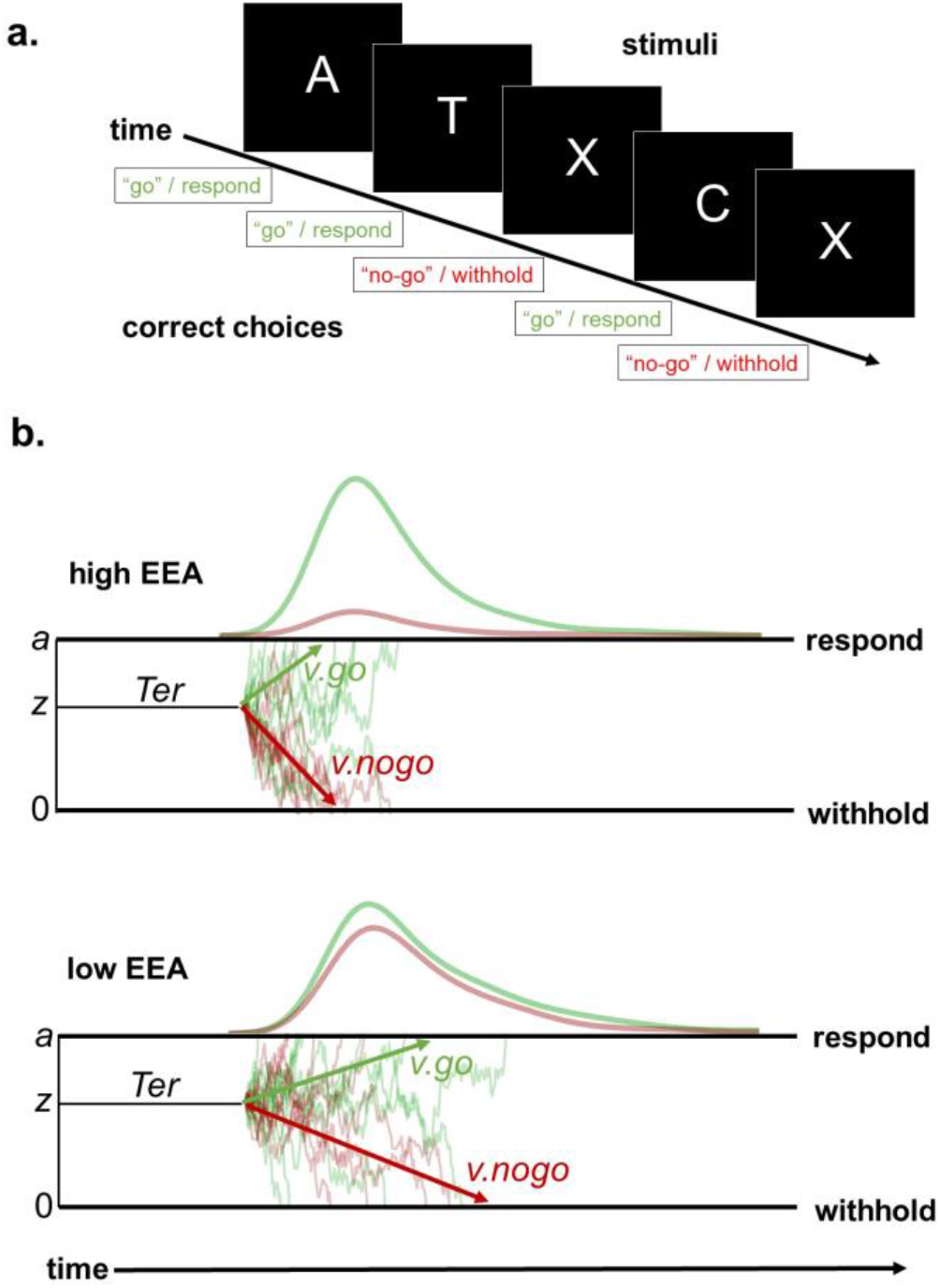
(**a**) Schematic of the go/no-go task. Participants are presented with a string of letters and are instructed to respond on trials with any letter other than “X” (“go” trials) but to withhold from responding on trials where an “X” is presented (“no-go” trials). (**b**) Simplified schematic of the diffusion decision model (DDM) description of the go/no-go task in cases with relatively high (top) vs. low (bottom) efficiency of evidence accumulation (EEA). The DDM assumes that responses on choice response time (RT) and go/no-go tasks are the result of a decision variable that drifts over time, on the basis of noisy evidence gathered from the stimulus, until it reaches one of two boundaries which represent each possible choice (e.g., to respond vs. withhold). When decision processes on individual “go” and “no-go” trials, which are represented by the light green and light red traces, respectively, terminate at one of the boundaries, the corresponding choice is made. The boundaries are set at 0 and parameter *a*, and the decision variable begins at a starting value *z*. A non-decision time (*Ter*) parameter accounts for time taken up by processes peripheral to the decision (e.g., the motor response). The drift rate parameter (*v*) determines the average rate at which the decision variable drifts towards the boundary for the correct choice (*v.go* for “go” trials, *v.nogo* for “no-go trials) and can be used as a measure of EEA in individual differences analyses. Lines above the “respond” boundaries represent RT distributions for responses on correct “go” trials (green) and failed “no-go” trials (red). The case with lower EEA exhibits fewer correct “go” responses, more incorrect “no-go” responses, and more variable RTs (a greater proportion of long RTs in the skewed right tail) due to lower *v*.

However, none of these longitudinal neuroimaging studies found associations between false alarm (FA) rate on “no-go” trials, the primary behavioral index of inhibition on the go/no-go task, and substance use, suggesting that inhibitory functioning may not be a straightforward predictor of future use. Furthermore, as none of these studies assessed prospective predictors of substance use in participants older than age 21, it remains unclear whether measures from the go/no-go predict use in individuals’ early-to-mid 20s, when SUDs most commonly emerge.

A broader limitation of this previous work relates to the construct of “inhibitory control” as an individual difference variable. Although inhibitory control and related “executive” constructs were validated with earlier factor analytic work (Miyake et al., 2000), more recent studies have provided compelling evidence that these constructs lack coherence as individual difference dimensions (Eisenberg et al., 2019; Karr et al., 2018). Moreover, because these cognitive functions were defined primarily based on behavioral test score covariation, rather than on an understanding of the mechanistic processes involved in cognition, they are likely to display only tenuous links to neural circuitry (Wiecki et al., 2015). Computational psychiatry, an emerging field that seeks to identify candidate biobehavioral dimensions that confer risk for psychopathology by focusing on formal model parameters that index mechanistic, biologically plausible processes underlying task behavior (Wang & Krystal, 2014; Montague et al., 2012; Huys et al., 2016; Adams et al., 2016), offers a promising alternative framework.

Indeed, the diffusion decision model (DDM) (Ratcliff, 1978; Ratcliff et al., 2016), which provides a quantitative description of how individuals perform the go/no-go task (Figure 1b), suggests at least one such dimension that may be highly relevant to substance use risk. The DDM explains behaviors on the go/no-go and other choice tasks as the result of a process in which a decision variable drifts over time, in a noisy pattern influenced by stochastic evidence accumulated from the stimulus, between two boundaries representing each possible choice. When the process terminates at one of these boundaries (e.g., the “respond” boundary in a go/no-go task), the corresponding choice is made. Crucially, the parameters posited by the DDM and related models align remarkably well with neurophysiological data (Cassey et al., 2016; Gold & Shadlen, 2007; Smith & Ratcliff, 2004). The critical “drift rate” (*v*) parameter, which determines the rate at which the decision variable drifts toward the boundary for the correct choice, is typically used in experimental work to index the quality of evidence that can be extracted from a stimulus (Ratcliff et al., 2016); stimuli that provide more ambiguous information about which choice is correct lead to relatively slower drift rates (reflecting poorer evidence quality).

When considered as an individual differences dimension, drift rate (*v*) indexes the efficiency with which individuals can gather relevant evidence to make an accurate choice in the context of background noise, or “efficiency of evidence accumulation” (EEA; Figure 1b). Measures of EEA display clear trait-like properties even when indexed across an array of simple and complex tasks from diverse cognitive domains (Lerche et al., 2020; Schubert et al., 2016). Furthermore, there is evidence that EEA may underlie performance on tasks thought to measure inhibition and other “executive” constructs (Evans et al., 2018; Karalunas & Huang-Pollock, 2013; Schmiedek et al., 2007; Weigard & Huang-Pollock, 2017), including the go/no-go (Huang-Pollock et al., 2017). We recently found (Weigard et al., 2019) that EEA on the go/no-go task, measured across conditions that both did (“no-go” trials) and did not (“go” trials) require inhibition, was strongly related to the inhibitory performance (FAs) of young adults and was robustly positively correlated with error-related activation in the anterior cingulate cortex (ACC) and anterior insula. Both regions are considered key hubs of the salience network (Seeley et al., 2007), a brain network involved in triggering cognitive control in response to errors and other salient events. These findings establish that EEA is a biobehavioral dimension that both drives individual differences in go/no-go task performance and has clear neuroimaging correlates.

As lower EEA has been repeatedly linked to externalizing psychopathologies comorbid with substance use (Endres et al., 2014; Weigard et al., 2018; Ziegler et al., 2016), previous links between go/no-go measures and use could be explained by reduced EEA rather than by selective deficits in inhibition. Crucially, this hypothesis differs from traditional theories of “inhibitory control” as a risk factor for substance use because it assumes EEA is a task-general neurocognitive factor, and should therefore relate to use regardless of whether EEA is measured under conditions that require inhibition (e.g., “no-go” trials) or those that do not (“go” trials).

In the current study, we build on our prior cross-sectional work (Weigard et al., 2019) to explicitly test, in longitudinal data, whether EEA and its neural correlates (error-related activations) prospectively predict individuals’ substance use during emerging adulthood. We tested two main hypotheses. First, on the basis of our previous work (Weigard et al., 2019), we expected that EEA measured on the go/no-go task would display a robust, positive relationship with a multivariate neural measure of error-related activation. Second, we predicted that both EEA and this error-related activation measure would display negative relationships with prospective substance use. Importantly, since we posit that EEA underlies performance in conditions that both do, and do not, require inhibition, we also expected that, for both of these hypothesized relationships, we would find similar associations when EEA is measured with: i) “go” trials; ii) “no-go” trials; and iii) a composite variable drawn from both types of trials.

## 2. Materials and Methods

### 2.1 Participants

Participants were volunteers for a functional magnetic resonance imaging (fMRI) substudy that was part of the Michigan Longitudinal Study (MLS) (Zucker et al., 1996, 2002), a prospective study that has followed a sample of youth from families with history of alcohol use disorder (AUD) and youth from low-risk families who lived in the same neighborhoods. MLS assessments began at ages 3–5 and were continued through participants’ late 20s and early 30s. Participants were excluded from the larger MLS if they displayed evidence of fetal alcohol syndrome and from the neuroimaging substudy if they: 1) had contraindications to MRI, 2) were left-handed, 3) had a neurological, acute, or chronic medical illness, 4) had a history of psychosis or first-degree relative with psychosis, or 5) were prescribed psychoactive medications, except psychostimulants prescribed for attention difficulties. Participants taking psychostimulants were asked to abstain from taking their medication for at least 48 hours prior to MRI scanning. Study procedures were carried out in accordance with the Declaration of Helsinki. Informed consent was obtained from all adult participants; for MLS waves in which participants were minors, parental consent and child assent were obtained.

For the current study, we examined MLS participants (*N*=143) who were included in our previous investigation of DDM parameters’ neural correlates from common go/no-go neuroimaging contrasts (Weigard et al., 2019). Individuals in this sample met all inclusion criteria outlined above and also had go/no-go behavioral and neuroimaging data available from their baseline scanning session (conducted at ages 18–21) that met quality-control criteria for DDM fitting and fMRI analysis (described in more detail in: Weigard et al., 2019). Of this sample, a subset of individuals (*n*=106; 74%) had substance use outcome data from at least one time point between ages 22 and 26. To optimize our ability to identify reliable metrics of error-related activation and link them to EEA, we used data from the *full sample* (*N*=143) for dimension reduction of fMRI data and assessment of whether our summary measure of error-related activation was related to drift rate. We then used data from the *longitudinal subsample* (*n*=106) to evaluate prospective relationships with substance use. Demographics and summary statistics of relevant variables for the full sample and longitudinal subsample are displayed in Table 1.

**Table 1.** Demographic information and summary statistics (means with standard deviations in parentheses) of all DDM parameters and variables included in prediction analyses. Demographics and statistics are reported separately for both the full sample included in the PCA of neural activation (*N*=143) and the subsample with available substance use outcome data (*n*=106), which was the focus of predictive analyses. “Annual” substance use measures were averaged over all available assessments from ages 22–26, and the “# of Measurements” rows report the mean number of measurement time points available per person for each substance use variable. AUD FHx = family history of alcohol use disorder (either parent); ADHD Dx = attention-deficit/hyperactivity disorder diagnosis; Cu. = cumulative sum of each substance use measure up to and including participants’ assessment at age 17; FA = false alarm

| Demographics/Measures | Full Sample<br>($N=143$) | Longitudinal<br>Subsample ($n=106$) |
| --- | --- | --- |
| Sex (Male/Female) | 87/56 | 63/43 |
| Age at scan | 19.66 (1.22) | 19.89 (1.20) |
| Race/Ethnicity |  |  |
| Caucasian | 128 | 102 |
| Hispanic/Latino | 5 | 2 |
| African American | 6 | 2 |
| Other/bi-racial | 4 | 0 |
| AUD FHx (Positive/Negative/Unknown) | 108/34/1 | 78/28/0 |
| ADHD Dx | 15 | 7 |
| FA rate | 0.28 (0.16) | 0.25 (.15) |
| Drift rate for “go” stimuli (*v.go*) | 2.77 (1.03) | 2.95 (1.03) |
| Drift rate for “no-go” stimuli (*v.nogo*)* | 1.90 (0.88) | 2.04 (0.80) |
| Average drift rate for all stimuli (*v.avg*) | 2.34 (0.88) | 2.50 (0.86) |
| Boundary separation (*a*) | 1.00 (0.21) | 1.00 (0.21) |
| Non-decision time (*Ter*) | .317 (.032) | .314 (.031) |
| Response bias (*z*) | 0.62 (0.07) | 0.62 (0.07) |
| Annual Drink Volume (ages 22–26) | --- | 429.91 (451.68) |
| Annual Marijuana Frequency (ages 22–26) | --- | 38.98 (83.70) |
| Annual Cigarette Frequency (ages 22–26) | --- | 89.14 (136.08) |
| # of Drink Volume Measurements | --- | 4.00 (1.32) |
| # of Marijuana Frequency Measurements | --- | 4.01 (1.33) |
| # of Cigarette Frequency Measurements | --- | 4.03 (1.33) |
| Cu. Drink Volume (at age 17) | 312.86 (781.19) | 381.70 (764.87) |
| Cu. Marijuana Frequency (at age 17) | 60.99 (178.93) | 48.38 (130.87) |
| Cu. Cigarette Frequency (at age 17) | 145.55 (356.51) | 147.26 (351.81) |

Descriptions of the go/no-go task completed by participants (which involved 245 trials, 60 of which were “no-go” trials), fMRI scanning parameters, and the fMRI pre-processing and single-subject analysis steps used in this and the previous study (Weigard et al., 2019) are provided in Supplemental Materials.

### 2.2 EEA Measure

As detailed in our prior study (Weigard et al., 2019), DDM parameters were estimated following methods established in a previous extension of the DDM to the go/no-go task (Huang-Pollock et al., 2017; Ratcliff et al., 2018) using functions from the R package *rtdists* (Singmann et al., 2016). Simplified versions of the DDM, without between-trial variability parameters, have been demonstrated to provide similar inferences to those drawn from the full DDM and may improve recovery of the main DDM parameters (Dutilh et al., 2019; Voss et al., 2013).

Therefore, we fit a version of the DDM that included only the following parameters: drift rates for “go” (*v.go*) and “no-go” (*v.nogo*) trials, starting point (*z*), boundary separation (*a*), and non-decision time (*Ter*). The simple average of drift rates across go and no-go trials (*v.avg*) provided a composite index of EEA. Additional information on model fit and parameter recovery studies is summarized in Supplemental Materials and provided at length in our previous study (Weigard et al., 2019).

### 2.3 Error-related Neural Activity Measure

We chose to use a multivariate measure of error-related activation because of recent indications that neural activity related to cognitive states and behavioral covariates is typically distributed across large brain networks, rather than being associated with discrete regions (Cohen et al., 2017; Yoo et al., 2019). We included regions of interest (ROIs) from our previous study (Weigard et al., 2019), obtained by conducting a term-based meta-analysis in Neurosynth (Yarkoni et al., 2016) with the term “error.” We downloaded the “association test” statistical map from this meta-analysis, which indicates whether activation in each voxel occurs more consistently in studies that mention the term “error” than in those that do not (see: neurosynth.org/faq). We excluded clusters with <10 voxels from this map (14 clusters between 2 and 6 voxels; 32 individual/isolated voxels) and then, for the 8 clusters that remained (minimum size = 28 voxels), centered 8mm radius spheres about their peak coordinates (Figure 2). Regions included ACC and two clusters spanning bilateral insula and nearby inferior frontal gyrus (IFG).

**Fig. 2.**
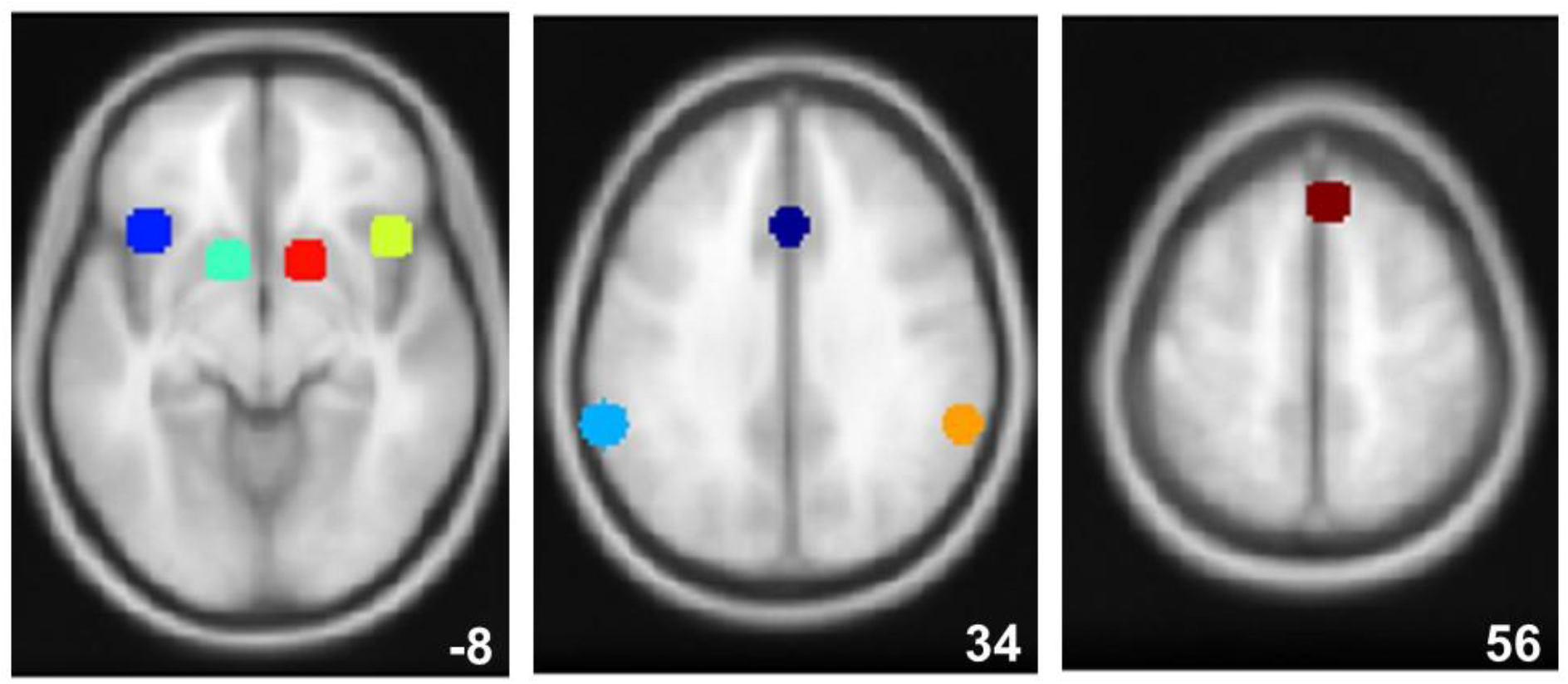
Spheres (8mm radius) centered about coordinates (see Table 3 in Results) of our 8 regions of interest (ROIs; white numbers in lower right indicate z-coordinates).

We next extracted average activation parameter estimates from the primary “error monitoring” contrast of interest (failed inhibitions on “no-go” trials > “go” responses) within each ROI, and entered these estimates into a principal component analysis (PCA) using the R package *FactoMineR* (Lê et al., 2008). We used scores of the first component (PC1) from this PCA as our primary measure of error-related activation in order to harness the aforementioned advantages of a network-based approach, and we anticipated that this component would display strong loadings from the ROIs closely associated with the salience network in previous literature (ACC and bilateral insula/IFG) (Seeley et al., 2007). However, for comprehensiveness, we also report prospective relationships between neural activations from discrete brain regions and substance use in Supplemental Materials. Furthermore, as results of the PCA also revealed a second component (PC2) that was of potential interest because of its links to striatal activation (see below), associations of this component with EEA and substance use were also assessed.

### 2.4 Substance Use Measures and Covariates

Substance use was assessed annually in the MLS sample with the Drinking and Drug History Form (Zucker et al., 1990). We aimed to create an outcome measure that indexed the degree to which individuals used the three most common substances of abuse—alcohol, marijuana and tobacco—during emerging adulthood, and therefore focused on three measures from this questionnaire: drink volume (number of alcoholic drinks in the past year), marijuana frequency (number of days in the past year when marijuana was used), and cigarette frequency (number of days in the past year when cigarettes were used). To obtain a stable measure of how frequently individuals used these substances during ages 22–26, participants’ responses for all ages with available data during this period were averaged for each substance (mean number of time points per measure are reported in Table 2). These three measures of average annual use during ages 22–26, which were moderately correlated with one another (Supplemental Table 1), were converted to Z-scores and averaged to form a substance use composite measure (SC) that was the main outcome measure of interest.

**Table 2.** Posterior median *r* values and Bayesian credible intervals (CIs) for correlations between the primary behavioral inhibition measure from the go/no-go task (FA rate) and drift rate measures. Values below the diagonal indicate relationships within the full sample (*N*=143; as previously reported in: Weigard et al., 2019) while those above the diagonal indicate relationships within the longitudinal subsample only (*n*=106). *v.go* = drift rate from “go” trials; *v.nogo* = drift rate from “no-go” trials; *v.avg* = composite of drift rate across all trials

|  | **FA rate** | ***v.go*** | ***v.nogo*** | ***v.avg*** |
| --- | --- | --- | --- | --- |
| FA rate | — | -0.73 | -.63 | -.73 |
|  | — | -0.62 | -.49 | -.62 |
|  | — | -0.80 | -.72 | -.80 |
| *v.go* | -.67 | — | .75 | .95 |
|  | -.56 | — | .82 | .97 |
|  | -.75 | — | .64 | .93 |
| *v.nogo* | -.66 | .73 | — | .92 |
|  | -.55 | .79 | — | .94 |
|  | -.74 | .63 | — | .88 |
| *v.avg* | -.71 | .94 | .92 | — |
|  | -.62 | .96 | .94 | — |
|  | -.78 | .92 | .88 | — |

Several covariates were used in predictive analyses to adjust for effects of other factors previously found to be related to substance use, including participants’ sex (0 = male, 1 = female), race/ethnicity (due to the small number of non-white participants: 1 = white, 0 = non-white), family history of AUD (given the enrichment of this sample with individuals who had this risk factor; 0 = no family history, 1 = AUD history for one or both parents), and participants’ level of substance use prior to the time of the scan (described in detail below). As a subset of individuals were diagnosed with attention-deficit/hyperactivity disorder (ADHD; Table 1), and as individuals taking medications for ADHD were asked to cease taking their medication >48 hours prior to scanning, we also included ADHD diagnosis (0 = negative, 1 = positive) as a covariate to account for possible confounds related to the disorder or to medication withdrawala. Although we considered including participants’ age at the scan as a covariate, we decided against this because of the narrow age range of the sample (18–21) and the fact that variability in age within this range was unrelated to either EEA (*v.avg*: *r*=.08, *p*=.40, BF10=0.17) or error-related activation (PC1: *r*=.00, *p*=.99, BF10=0.12).

Prior substance use covariates were cumulative sums of the same three common substance use measures (drink volume, marijuana frequency, and cigarette frequency) from annual assessments up to and including individuals’ assessment at age 17. Similar to our outcome measure, we converted these covariates to Z-scores and averaged them to form a prior substance use composite measure (preSC), which was used as a single covariate in regression models. However, we also conducted a sensitivity analysis (Supplemental Materials) in which raw scores of individual prior substance use covariates were used in place of preSC. Notably, preSC did not show evidence of a relationship with either EEA (*v.avg*: *r*=−.09, *p*=.37, BF10=0.18) or error-related activation (PC1: *r*=.07, *p*=.47, BF10=0.16), suggesting that prior substance exposure does not have a substantial impact on individuals’ EEA in this age range.

### 2.5 Inferential Analyses

We conducted all inferential analyses using JASP (JASP Team, 2020), a software package allowing users to conduct frequentist and Bayesian statistical tests. We conducted analyses from both statistical frameworks in order to evaluate whether our results were sensitive to the chosen framework and to leverage Bayesian model comparison methods to obtain continuous estimates of the probability that our two primary variables of interest, the EEA composite (*v.avg*) and error-related activation, provided unique predictive information.

First, we used Pearson correlation (*r*) tests to assess whether EEA on the behavioral task, estimated across conditions that both did and did not require inhibition (*v.go*, *v.nogo*) and using the composite (*v.avg*), displayed a positive relation with error-related activation (PC1). These inferences were evaluated with both frequentist *p*-values, which test whether the null hypothesis (*r*=0) can be rejected, and Bayes factors (BF), which are continuous measures of evidence that the data provide for an alternative model/hypothesis (H1) relative to the null model/hypothesis (H0). The BF10 is intuitively interpreted as an odds ratio; a BF10 of 6.00 in favor of H1 indicates that the data are 6 times more likely under H1 than H0, while a BF10 of 0.33 would indicate that the data are instead BF01=3 (=1/0.33) times more likely under H0 than H1. In Bayesian correlation tests, H1 and H0 correspond to the hypotheses that *r*≠0 and *r*=0, respectively^b^.

Second, frequentist regression analyses were conducted in which all measures of EEA (*v.go*, *v.nogo*, *v.avg*), error-related activation (PC1), and the striatal activation component (PC2) were separately entered along with covariates as possible predictors of the SC measure. These analyses tested whether the null hypothesis (i.e., our covariates of interest are not prospectively related to SC when accounting for other previously established risk factors) could be rejected in each case. We used false discovery rate (FDR: *q*<.05), where each separate regression was considered its own family of tests, to assess whether *p*-values from this analysis were robust to correction for multiple comparisons. For comparison, we also entered the primary behavioral index of inhibition from the go/no-go task (FA rate) into similar analyses in Supplemental Materials.

Finally, we conducted Bayesian linear regression analyses (Li & Clyde, 2018; Liang et al., 2008; Rouder & Morey, 2012). To study the relationships between the main covariates of interest, EEA and error-related activation, and the outcome SC after accounting for effects of covariates, we first added all nuisance covariates (e.g., preSC) to a “null” model. We then estimated alternative models of interest, which included these nuisance covariates as well as all possible combinations of the covariates of interestc. As EEA measures across conditions were very strongly correlated (see below), precluding their simultaneous inclusion in regression analyses, and as cross-condition EEA was our main covariate of interest for the theoretical reasons outlined above, only *v.avg* and PC1 were considered as covariates of interest. Models were then compared using BF10, which quantifies the evidence in favor of each alternative model relative to the “null” model, and BFM, which quantifies evidence for each model compared to all other models. To summarize the importance of EEA and error-related activation across all models, we also performed model averaging, which provides us with evidence for inclusion, relative to non-inclusion of each variable (BF_inc_) (Clyde et al., 2011; Ly et al., 2019).

Importantly, models that are averaged in this way provide evidence that is corrected for multiple testing (Scott et al., 2010; van den Bergh et al., in press). These Bayesian analyses allowed us to directly test whether *EEA* and error-related activation contain unique versus redundant predictive information by quantifying evidence for a model that contains both variables relative to simpler models that contain only one of each.

## 3. Results

### 3.1 Drift Rate and Behavioral Inhibition Measures

Table 2 displays correlations between the main behavioral index of inhibition on the task (FA rate) and measures of EEA in both the full sample and the longitudinal subsample. As we previously reported, FA rate is strongly correlated with EEA, regardless of whether EEA is measured on “go” trials (*v.go*), “no-go” trials (*v.nogo*), or with the cross-condition EEA composite (*v.avg*). Combined with the fact that *v.go* and *v.nogo* are also highly intercorrelated, this finding suggests it is well-justified to view EEA on the task as a single individual difference dimension that similarly determines both inhibitory performance and performance in the “go” condition. Furthermore, it raises practical considerations about including FA rate and the three EEA measures in the same regression analyses; as these measures explain roughly 50% of the variance in one another, such regression models are unlikely to be informative due to multicollinearity problems.

### 3.2 Summary of Error-related Activation

Results from the PCA of error-related activation in our 8 ROIs are displayed in Table 3. Five components were necessary to explain more than 90% of the variance, although the first component explained the majority (51.22%). As expected, the first component was highly correlated with error-related activation in all ROIs, and salience network structures displayed particularly strong loadings on this component, including ACC (.81), right IFG/insula (.78), and left IFG/insula (.81). Notably, striatal ROIs displayed the lowest loadings on the first component (.45 and .55), and these ROIs were, instead, selectively related to the second component (PC2). This pattern may reflect the fact that, although bilateral striatum was identified in our term-based meta-analysis as being related to the term “error,” these structures are not typically associated with error monitoring or the salience network. Their identification in the meta-analysis may have been an artifact of the inclusion of the word “error” in other constructs associated with striatum (e.g., “reward prediction error”) (Abler et al., 2006). As error-related activations in ROIs previously associated with error monitoring and the salience network were strongly related to PC1, we concluded that this component would operate as an effective summary measure of error-related activation. However, as the striatum-specific component (PC2) may be of interest as well, we also assessed this component’s associations with EEA and substance use in subsequent analyses.

**Table 3.**
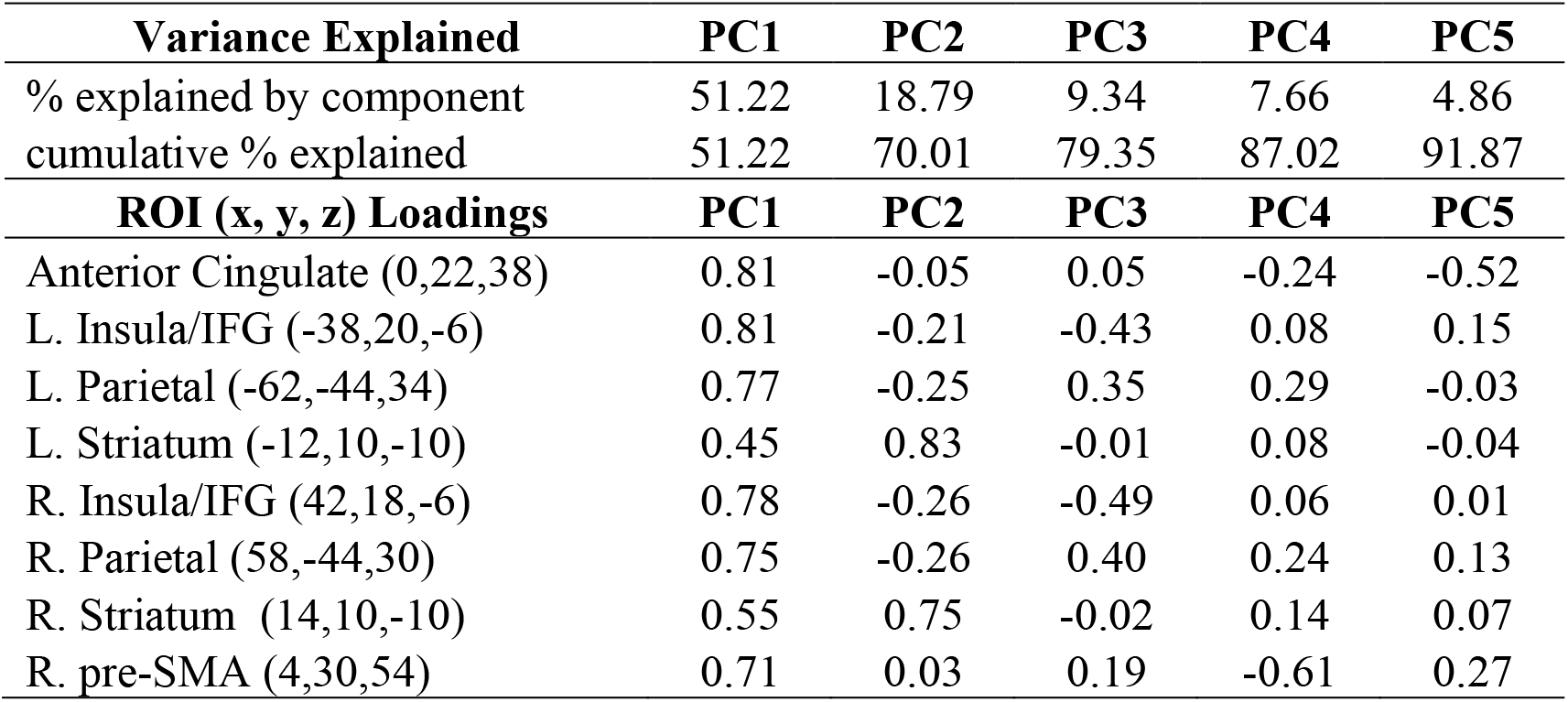
Variance explained by, and loadings of each region of interest (ROI) on, the five principal components (PCs) that together explain over 90% of between-subject variation in error-related ROI activations. MNI coordinates of ROIs are displayed in parentheses. L. = left; R. = right; IFG = inferior frontal gyrus; SMA = supplementary motor area

| **Variance Explained** | **PC1** | **PC2** | **PC3** | **PC4** | **PC5** |
| --- | --- | --- | --- | --- | --- |
| % explained by component | 51.22 | 18.79 | 9.34 | 7.66 | 4.86 |
| cumulative % explained | 51.22 | 70.01 | 79.35 | 87.02 | 91.87 |
| **ROI (x, y, z) Loadings** | **PC1** | **PC2** | **PC3** | **PC4** | **PC5** |
| Anterior Cingulate (0,22,38) | 0.81 | -0.05 | 0.05 | -0.24 | -0.52 |
| L. Insula/IFG (-38,20,-6) | 0.81 | -0.21 | -0.43 | 0.08 | 0.15 |
| L. Parietal (-62,-44,34) | 0.77 | -0.25 | 0.35 | 0.29 | -0.03 |
| L. Striatum (-12,10,-10) | 0.45 | 0.83 | -0.01 | 0.08 | -0.04 |
| R. Insula/IFG (42,18,-6) | 0.78 | -0.26 | -0.49 | 0.06 | 0.01 |
| R. Parietal (58,-44,30) | 0.75 | -0.26 | 0.40 | 0.24 | 0.13 |
| R. Striatum (14,10,-10) | 0.55 | 0.75 | -0.02 | 0.14 | 0.07 |
| R. pre-SMA (4,30,54) | 0.71 | 0.03 | 0.19 | -0.61 | 0.27 |

### 3.3 Neural Correlates of EEA

There was a robust positive correlation between PC1 and all three indices of EEA (Figure 3a), both in the full sample (*v.go r*=.30, *p*<.001, BF10=62.48; *v.nogo r*=.30, *p*<.001, BF10=77.89; *v.avg r*=.32, *p*<.001, BF10=204.43), and when the longitudinal subsample was considered separately (*v.go r*=.23, *p*=.017, BF10=2.03; *v.nogo r*=.23, *p*=.02, BF10=1.78; *v.avg r*=.25, *p*=.011, BF10=2.87). In contrast, the correlational relationship between PC2 and EEA measures, both in the full sample (*v.go r*=-.13, *p*=.125, BF10=0.34; *v.nogo r*=−.14, *p*=.092, BF10=0.43; *v.avg r*=−.15, *p*=.084, BF10=0.46) and longitudinal subsample (*v.go r*=−.13, *p*=.198, BF10=0.28; *v.nogo r*=−.15, *p*=.128, BF10=0.38; *v.avg r*=−.15, *p*=.137, BF10=0.36) was systematically nonsignificant. Hence, error-related activation from the salience network component (PC1) displayed consistent evidence of a positive relationship with EEA measured across both “go” and “no-go” conditions, as expected.

**Fig. 3.**
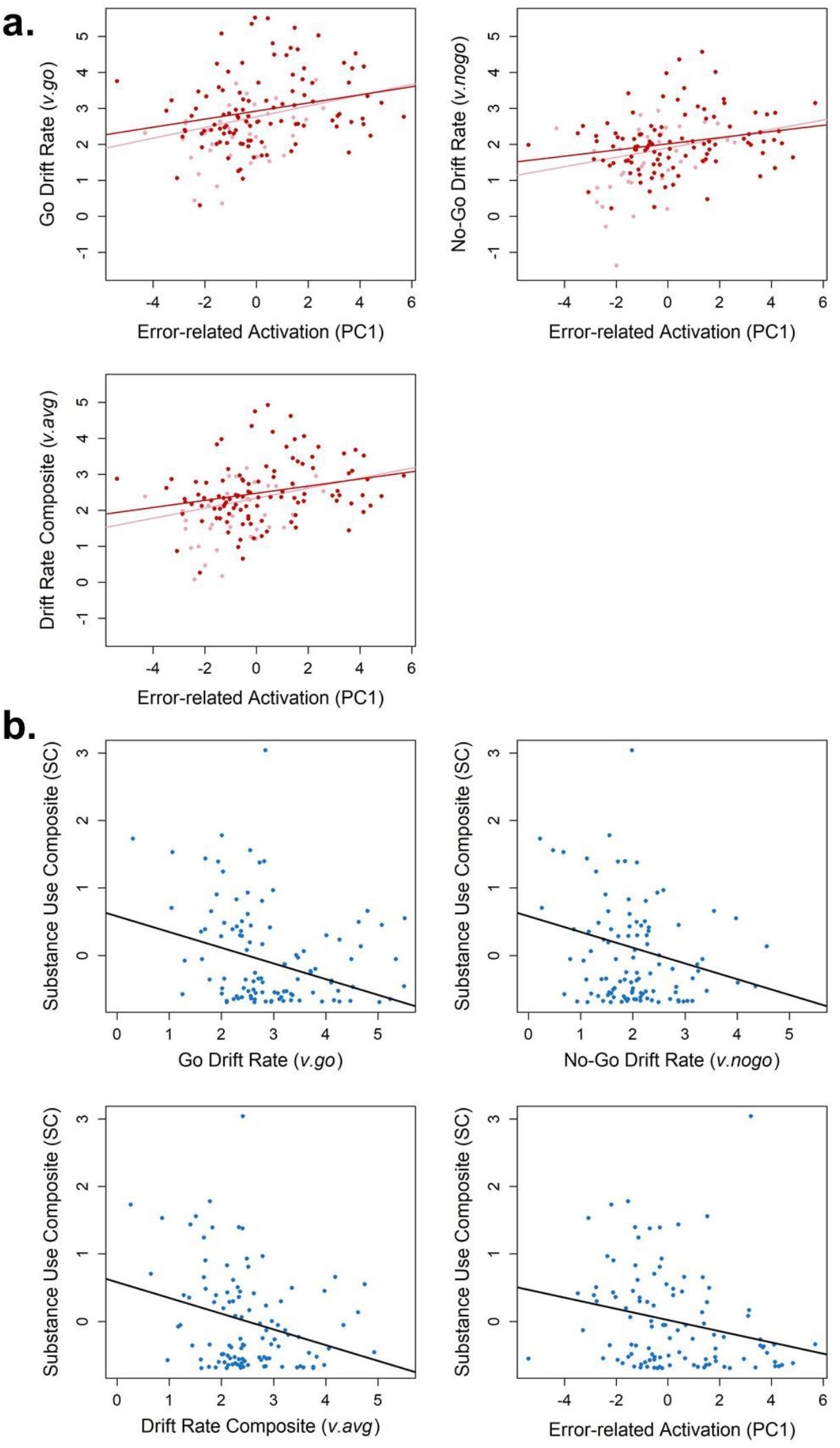
(**a**) Scatterplots and simple regression lines of the association between our summary measure of error-related activation (PC1) and individuals’ drift rates from the go/no-go task, including drift rate on “go” trials (*v.go*), drift rate on “no-go” trials (*v.nogo*), and the composite drift rate measure from across all trials (*v.avg*). Red points and lines represent these relationships for the subsample of individuals with substance use outcome data (*n*=106) and pink points and lines represent the same relationships when all individuals in the full sample (*N*=143) are included. (**b**) Scatterplots of associations in which drift rates (*v.go*, *v.nogo*, *v.avg*) and our summary measure of error-related activation (PC1) predict individuals’ values of the age 22–26 substance use composite (SC). Simple regression lines are displayed in black.

### 3.4 Frequentist Regression Analyses

In frequentist regression analyses considering each covariate of interest along with other possible risk factors for substance use (Table 4; scatterplots in Figure 3b), *v.go*, β=−0.23, *p*=.006, *v.avg*, β=−0.21, *p*=.011, and PC1, β=−0.25, *p*=.002, were all found to have statistically significant negative associations with the SC from ages 22–26, but *v.nogo*, β=−0.16, *p*=.059, and PC2, β=0.11, *p*=.226, were not. Male sex and, even more so, greater prior substance use also appeared to be predictors of the SC across regressions, which was unsurprising considering previous research on these factors. The behavioral measure of inhibition (FA rate) did not display a significant association with the SC, β=0.15, *p*=.088 (Supplemental Materials), consistent with previous findings (Heitzeg et al., 2014; Mahmood et al., 2013; Wetherill et al., 2013). Notably, although *v.nogo* did not reach significance as a predictor, it displayed a qualitative relationship with the SC that was highly similar to that of the other EEA measures (Figure 3b). Overall, these analyses suggest that substance use is prospectively predicted by both lower levels of error-related activation in salience network regions during the task (PC1) and lower cross-condition EEA (*v.avg*), although EEA measured on “go” trials appears to be a more robust predictor than EEA on “no-go” trials.

**Table 4.** Results from frequentist regression analyses predicting values of the age 22–26 substance use composite (SC) with models that included measures of EEA (*v.go*, *v.nogo*, *v.avg*), PC1, and PC2, along with covariates. **Bolded***p*-values survive false discovery rate correction for multiple comparisons within families defined by the individual regression models. Overall variance explained by each model (R^2^) is displayed in parentheses. R/E = race/ethnicity; AUD FHx = family history of alcohol use disorder (either parent); ADHD Dx = attention-deficit/hyperactivity disorder diagnosis; preSC = prior substance use composite (cumulative use through age 17)

| Model<br>($R^2$) | | Unstandardized | Standard<br>Error | Standardized | *T* | *p* |
| --- | --- | --- | --- | --- | --- | --- |
| ***v.go***<br>**(.38)** | (Intercept) | 0.469 | 0.358 |  |  |  |
|  | Sex | -0.297 | 0.117 | -0.202 | -2.536 | **0.013** |
|  | R/E | 0.071 | 0.302 | 0.019 | 0.236 | 0.814 |
|  | AUD FHx | 0.043 | 0.132 | 0.026 | 0.323 | 0.747 |
|  | ADHD Dx | 0.326 | 0.234 | 0.112 | 1.394 | 0.166 |
|  | preSC | 0.405 | 0.065 | 0.496 | 6.217 | **< .001** |
|  | *v.go* | -0.159 | 0.057 | -0.227 | -2.790 | **0.006** |
| ***v.nogo***<br>**(.36)** | (Intercept) | 0.32 | 0.364 |  |  |  |
|  | Sex | -0.273 | 0.12 | -0.186 | -2.28 | 0.025 |
|  | R/E | 0.049 | 0.31 | 0.013 | 0.159 | 0.874 |
|  | AUD FHx | 0.021 | 0.135 | 0.013 | 0.157 | 0.876 |
|  | ADHD Dx | 0.337 | 0.239 | 0.116 | 1.41 | 0.162 |
|  | preSC | 0.417 | 0.066 | 0.510 | 6.288 | **< .001** |
|  | *v.nogo* | -0.144 | 0.076 | -0.160 | -1.911 | 0.059 |
| ***v.avg***<br>**(.38)** | (Intercept) | 0.466 | 0.365 |  |  |  |
|  | Sex | -0.283 | 0.118 | -0.192 | -2.401 | **0.018** |
|  | R/E | 0.047 | 0.304 | 0.012 | 0.156 | 0.877 |
|  | AUD FHx | 0.041 | 0.133 | 0.025 | 0.305 | 0.761 |
|  | ADHD Dx | 0.319 | 0.236 | 0.110 | 1.355 | 0.179 |
|  | preSC | 0.409 | 0.065 | 0.501 | 6.253 | **< .001** |
|  | *v.avg* | -0.179 | 0.069 | -0.212 | -2.583 | **0.011** |
| **PC1**<br>**(.40)** | (Intercept) | -0.105 | 0.311 |  |  |  |
|  | Sex | -0.248 | 0.117 | -0.169 | -2.124 | 0.036 |
|  | R/E | 0.173 | 0.299 | 0.046 | 0.578 | 0.564 |
|  | AUD FHx | 0.055 | 0.131 | 0.033 | 0.416 | 0.678 |
|  | ADHD Dx | 0.306 | 0.232 | 0.105 | 1.319 | 0.19 |
|  | preSC | 0.436 | 0.064 | 0.534 | 6.77 | **< .001** |
|  | PC1 | -0.087 | 0.028 | -0.253 | -3.12 | **0.002** |
| **PC2**<br>**(.35)** | (Intercept) | -0.01 | 0.322 |  |  |  |
|  | Sex | -0.328 | 0.123 | -0.223 | -2.659 | **0.009** |
|  | R/E | 0.169 | 0.312 | 0.045 | 0.543 | 0.588 |
|  | AUD FHx | -0.054 | 0.137 | -0.033 | -0.392 | 0.696 |
|  | ADHD Dx | 0.363 | 0.241 | 0.125 | 1.505 | 0.135 |
|  | preSC | 0.424 | 0.067 | 0.519 | 6.342 | **< .001** |
|  | PC2 | 0.06 | 0.047 | 0.108 | 1.266 | 0.208 |

### 3.5 Bayesian Model Comparison

Due to our interest in cross-condition EEA, and the fact that PC2 did not appear to be a robust predictor of the SC in frequentist analyses, only the composite measure of EEA (*v.avg*) and PC1 were included as predictors of interest. Results from Bayesian regression analyses predicting the SC (Table 5) indicated that there was moderate to strong evidence for models that included *v.avg*, PC1, and both predictors simultaneously; BF10 indicated the observed data were 5.44, 19.72, and 35.10 times more likely under each of these models, respectively, than under the “null” (nuisance covariate only) model. BF10 values also indicated that the data were roughly 6.45 (35.10/5.44) times more likely under the model that simultaneously included *v.avg* and PC1 than under the model that included *v.avg* only, and 1.78 (35.10/19.72) times more likely under the two-predictor model than under the model that included PC1 only. Furthermore, BFM values, which quantify evidence for each model compared to all other models, indicated that the data provided moderate support for the model with both predictors of interest (BFM=4.03), comparatively modest support for the PC1-only model (BFM=1.42), and evidence against the *v.avg*-only model (BFM=0.29). In other words, BF10 and BFM both indicate that, although all models involving *v.avg* and PC1 are well-supported, there is some evidence that simultaneous inclusion of both predictors leads to a better description of the data than when only one is included.

**Table 5.** Results of Bayesian regression analyses in which all possible models involving the predictors of interest, cross-condition EEA (*v.avg*) and general error-related activation (PC1), were compared to a “null” model that included only the covariates of sex, race/ethnicity (R/E), family history of alcohol use disorder (AUD FHx), attention-deficit/hyperactivity disorder diagnosis (ADHD Dx), and the prior substance use composite (preSC). Note that all models contain these covariates in addition to any predictors of interest. In the “Model Comparison” section, P(M) is prior probability of the model, P(M|data) is the posterior probability of the model after seeing the data, BF10 is the Bayes factor comparing the model to the “null” model, and BFM is a Bayes factor comparing the model to all other models from the analysis. The “Posterior Summaries” section reports the model-averaged mean, standard deviation (SD) and 95% credible intervals of posterior samples for coefficients of each predictor of interest, as well as inclusion probabilities obtained from model averaging; P(inc) is the prior probability of including each predictor, P(inc|data) is the posterior probability of including each predictor, and BF_inc_ is a Bayes factor for the change from prior to posterior inclusion odds for the predictor after seeing the data.

| **Model Comparison** |  |  |  |  |  |  |  |
| --- | --- | --- | --- | --- | --- | --- | --- |
| **Models** |  |  | **P(M)** | **P(M\|data)** | **BF<sub>M</sub>** | **BF<sub>10</sub>** | **R<sup>2</sup>** |
| “Null” (Sex, R/E, AUD FHx, ADHD Dx, preSC) |  |  | 0.250 | 0.016 | 0.05 | 1.000 | 0.336 |
| *v.avg* + PC1 |  |  | 0.250 | 0.573 | 4.027 | 35.101 | 0.420 |
| PC1 |  |  | 0.250 | 0.322 | 1.424 | 19.715 | 0.396 |
| *v.avg* |  |  | 0.250 | 0.089 | 0.292 | 5.436 | 0.378 |
| **Posterior Summaries of Coefficients** |  |  |  |  |  |  |  |
| **Coefficient** | **Mean** | **SD** | **P(inc)** | **P(inc\|data)** | **BF<sub>inc</sub>** | **95% Credible Interval** |  |
|  |  |  |  |  |  | **Lower** | **Upper** |
| *v.avg* | -0.087 | 0.082 | 0.500 | 0.662 | 1.957 | -0.228 | 0.000 |
| PC1 | -0.064 | 0.034 | 0.500 | 0.895 | 8.517 | -0.116 | 0.000 |

Inclusion Bayes factors (BF_inc_) obtained via model averaging indicated positive evidence for inclusion of *v.avg*, BF_inc_=1.96, and PC1, BF_inc_=8.52, although evidence for the former was weaker. Finally, consideration of models’ explanatory power (*R*^2^) indicated that the model containing both *v.avg* and PC1 explained a substantially greater proportion of the variance than the “null” (covariate only) model (roughly 8% more). In sum, Bayesian analyses provide evidence that both *v.avg* and PC1 are meaningful predictors of substance use in emerging adulthood and that each may provide unique predictive information, even when included in the same model.

## 4. Discussion

This study aimed to test whether lower efficiency of evidence accumulation (EEA), a neurocognitive individual difference variable indexed by the DDM’s “drift rate” parameter (Ratcliff et al., 2016, 2018), is a risk factor for frequent substance use in emerging adulthood. In a longitudinal study, we found evidence that both lower levels of EEA as well as reductions in a robust neural-level correlate of EEA (error-related activation in brain regions linked to salience and performance monitoring) were prospectively related to greater use of the major substances of abuse (alcohol, marijuana, cigarettes) during ages 22–26.

Our prediction that EEA would be robustly related to substance use across trials that both did (“no-go”) and did not (“go”) require inhibition was only partially supported; we found that EEA on “go” trials and an EEA composite measure across both types of trials each significantly predicted use, while EEA on “no-go” trials only displayed a similar qualitative trend. However, when combined with evidence that EEA is likely a single individual difference dimension that determines performance across a variety of tasks and conditions, these findings are generally consistent with the idea that EEA displays a task-general predictive relationship with substance use. Indeed, it is plausible that EEA measured on “go” trials may be a more robust predictor of use than EEA on “no-go” trials simply due to reduced measurement error, since there are many more “go” trials compared to “no-go” trials on the task.

Our finding that EEA facilitates meaningful predictions about substance use in an age range critical for the emergence of SUDs has at least two major implications. First, although other cognitive constructs have been posited as risk factors for substance use problems (e.g., inhibitory control), these constructs have recently been criticized for lacking coherence as individual difference dimensions (Eisenberg et al., 2019; Karr et al., 2018) and lacking links to specific computational and neural mechanisms that can explain (rather than simply describe) cognitive performance (Wiecki et al., 2015). In contrast, EEA is a dimension derived from well-validated quantitative models that explain cognitive performance using formally specified and biologically plausible mechanisms (Cassey et al., 2016; Smith & Ratcliff, 2004), and EEA shows clear trait-like qualities when measured across a variety of tasks with different cognitive demands (Schmiedek et al., 2007; Schubert et al., 2016). Crucially, the latter implies that the relationships identified in the current study are likely not limited to the go/no-go or other “inhibition” tasks, and would be expected to be identified even when EEA is measured on relatively simple decision tasks that are not typically thought of as “executive” function measures (e.g., perceptual discrimination). If low EEA proves to be a task-general mechanism that underlies relationships between poorer performance on neurocognitive measures and later substance use problems, researchers could leverage these well-developed computational models, and knowledge of their links to neural processes (e.g., salience network activity), to identify circuits related to addiction risk.

Second, our findings indicate that EEA and its neural-level correlates may each provide unique information for substance use prediction, suggesting that inclusion of both measures in cross-validated models may ultimately enhance prediction of individual-level substance use outcomes in an applied context. Our Bayesian model comparison analyses provided moderate evidence that there is added value in including neural-level correlates of EEA in prediction models, even in addition to estimates of EEA itself. Hence, future work that seeks to utilize EEA to make real-world predictions about individuals’ substance use problems should consider identifying neural correlates of EEA that can be feasibly measured in applied settings (e.g., electrophysiological measures of error-monitoring). Additionally, as recent work suggests that self-report measures have greater predictive power than task-based measures (Eisenberg et al., 2019), identification of self-report measures that index dimensions similar to EEA may aid prediction. Future work establishing the place of EEA in a larger nomological network of task and survey measures is therefore needed.

The neural processes involved in evidence accumulation on choice task trials are well-understood at the computational and neurophysiological levels (Cassey et al., 2016; Smith & Ratcliff, 2004). However, investigations of systems-level correlates of individual differences in EEA are only getting started. Although EEA is likely to display multiple circuit- and systems-level neural correlates in different imaging modalities, the current study focused on error-related brain activation patterns because these were identified as robust neural correlates of EEA in our previous study (Weigard et al., 2019). Our finding that a summary measure of error-related activation was associated with EEA, and similarly predicted substance use outcomes, is significant in the context of accounts linking EEA to catecholamine systems thought to regulate arousal and optimize task performance in response to feedback (Aston-Jones & Cohen, 2005). Brain regions involved in performance monitoring are thought to provide input to these systems (Aston-Jones & Cohen, 2005), and connectivity of the salience network, comprised of regions that were major contributors to our summary error-related activation measure, has been specifically linked to norepinephrine action (Hermans et al., 2011). Therefore, individual differences in EEA may, in part, reflect individual differences in the integrity of catecholamine systems and associated neural networks that optimize task performance in response to external feedback or environmental demands.

This study has many strengths, but there are some limitations that should also be acknowledged. First, our sample was not large enough to use cross-validation methods to assess out-of-sample accuracy of our predictive models, which is necessary to provide accurate estimates of a model’s ability to predict new data in the real world. This is due to the fact that samples smaller than 150 subjects typically provide overly optimistic estimates of predictive accuracy due to model overfitting (Arbabshirani et al., 2017; Sui et al., 2020). Larger samples, such as that of the currently in-process Adolescent Brain Cognitive Development (ABCD) study (Casey et al., 2018), could be utilized in the future to assess whether measures of EEA, and its neuroimaging correlates, can predict substance use in unseen data. Second, we did not assess whether EEA shows a selective association with individual substances. We opted to predict a composite measure of common substance use because our sample was not large enough to identify predictors of rarer substances (e.g., opioids), or to identify distinctive predictors of the use of commonly used individual substances, measures of which were moderately correlated. Finally, although this study indicates that EEA and its neural correlates prospectively relate to substance use, the behavioral mediators by which low EEA confers risk are not currently known.

## 5. Conclusions

In sum, the current study provides evidence that lower levels of EEA, a biobehavioral dimension that is precisely measured in well-established computational models of brain function and has clear links to neuroimaging measures, shows promise as a risk factor for frequent substance use in emerging adulthood, a critical developmental period in the emergence of SUDs. These findings could inform predictive models of the emergence of substance use problems and take a crucial step in bringing the benefits of computational psychiatry to the developmental neuroscience of addiction.

## Acknowledgements

This project was supported by National Institute on Alcohol Abuse and Alcoholism (NIAAA) grants R01 AA07065 and R01 AA025790. Alexander Weigard and Sarah Brislin were supported by T32 AA007477. Jillian Hardee was supported by K01 AA024804. Lora Cope was supported by K01 DA044270. Meghan Martz was supported by K01 AA027558. Chandra Sripada was supported by R01 MH107741 and the Dana Foundation David Mahoney Neuroimaging Program. The authors have no conflicts of interest to declare.

## Author Contributions

RAZ and MH designed the larger study and acquired data for the data set that was utilized in the current work. AW, MH and CS conceptualized the research questions and analyses. AW analyzed and the data with assistance and consultation from AL and under the supervision of MH and CS. All authors were involved in interpretation of the analyses. AW, SB, MM, and LC wrote the first draft of the paper, and JH, AL, RAZ, CS and MH provided critical revisions. All authors approved the final version of the manuscript for submission and agree to be accountable for all aspects of the work. The authors have no competing interests to declare.

## Footnotes

a ADHD diagnosis was assessed with both the Diagnostic Interview Schedule (DIS) (Robins et al., 1981) and screening questions about previous ADHD diagnosis and medication use. We used diagnostic information from the DIS for all participants except for three who did not complete this measure. We therefore used information from the screening questions to assess diagnostic status for these three participants, all of whom reported no previous ADHD diagnosis or medication use. Given the small number of participants with ADHD in the longitudinal sample, it was not feasible to account for diagnosis and medication use with separate covariates.

b All correlation tests used JASP’s default prior on *ρ* is a uniform distribution spanning the values between −1 and 1 (Ly et al., 2016, 2018).

c All Bayesian regression analyses used a JZS prior with prior scale (0.354), which is the default prior in JASP.

