## Supplemental Materials for "Evidence accumulation and associated error-related brain activity as computationally-informed prospective predictors of substance use in emerging adulthood"

**Go/No-Go Task**

Participants completed an event-related go/no-go task (Durston et al., 2002; Heitzeg et al., 2014; Weigard et al., 2019) during fMRI data collection in which they were presented with a string of letters (white on black background) that indicated whether they should respond (any letter other than “X”; 75% of trials) or inhibit their response (“X”; 25% of trials). Letters were presented for 500ms (3500ms interstimulus fixation interval) during 5 imaging runs of 49 trials each (245 trials total; 60 “X” trials).

**MRI Data Acquisition Parameters**

A high-resolution T1-weighted anatomical image was obtained using the following parameters: three-dimensional spoiled gradient-recalled echo, TR=25ms, minimum TE, FOV=25cm, 256x256 matrix, slice thickness=1.4mm. During runs of the go/no-go task, whole brain T2*-weighted functional images were acquired using a single-shot spiral in-out sequence(Glover & Law, 2001) with the following parameters: TR=2000ms, TE=30ms, flip angle=90°, FOV=200mm, 29 axial slices, 64×64 matrix, in-plane resolution=3.12mm×3.12mm, and slice thickness=4mm. All scans were conducted with the same 3.0 T GE Signa scanner.

**Pre-Processing and Single-Subject fMRI Analyses**

Functional images were reconstructed using an iterative algorithm (Fessler et al., 2005) and entered into the following pre-processing steps: 1) motion correction with realignment using FSL 5.0.2.2 tools (FMRIB, Oxford, United Kingdom), 2) spatial normalization to standard space as defined by the Montreal Neurological Institute template using Statistical Parametric Mapping 8 (SPM8: Wellcome Institute of Cognitive Neurology, London, United Kingdom) and using normalization of the T1-weighted anatomical image for guidance, 3) resampling to 2x2x2mm voxels in SPM8, and 4) spatial smoothing with a 6mm full-width half-maximum Gaussian kernel. Functional runs were excluded from further analysis if they exceeded 3 mm translation or 3° rotation in any direction during the run.

A general linear model was fit in SPM8 to individual subjects’ fMRI time series data with three regressors convolved with the hemodynamic response function: 1) “go” responses, 2) successful inhibition (SI) trials, in which participants withheld their response to a “no-go” stimuli, and 3) failed inhibition (FI) trials, in which participants made a response following “no-go” stimuli. Motion parameters from earlier realignment and average white matter signal intensity for each volume were also included as nuisance regressors. Individual statistical maps for the primary “error monitoring” contrast of interest (FI > correct “go”) were generated for later analyses.

**Summary of Diffusion Decision Model Fits**

Detailed descriptions of how the diffusion decision model (DDM) (Ratcliff, 1978; Ratcliff et al., 2016) parameters were estimated, assessment of model fit, and assessment of parameter recovery, are available in our prior study (Weigard et al., 2019). Briefly, we followed the implementation of the DDM and chi-square minimization fitting routine that was outlined in two recent studies (Huang-Pollock et al., 2017; Ratcliff et al., 2018). Decisions to respond in the go/no-go were assumed to be made when the diffusion process crossed the upper response boundary while decisions to withhold from responding were assumed to be made when the process crossed the lower boundary. Therefore, when computing chi-square values for parameter estimation, upper boundary decisions (correct “go” responses and false alarms) were separated into 10 RT bins (denoted by quantiles .1 through .9) while lower boundary decisions (“go” omissions and correctly-inhibited “no-go” trials), which are not observed, were placed in a single “non-response” bin. RT quantiles were determined using functions from the R package *rtdists* (Singmann, Brown, Gretton, & Heathcote, 2016) and individuals’ parameters were estimated by minimizing the resulting chi-square value using the optim() function in R.

Model fit was assessed by plotting empirical accuracy rates and RT quantiles (.1, .5 and .9) from the “go” and “no-go” conditions against the same values predicted by the DDM. As described in detail in the Supplemental Material of our previous study (Weigard et al., 2019), the DDM provided an excellent description of accuracy rates and “go” trial RTs, but a poorer description of “no-go” trial RTs. Potentially because of the relative sparsity of data on “no-go”, relative to “go”, trials, there was more error in these predictions and an apparent bias in which the DDM over-predicted “no-go” trial RTs, especially for the longest RT quantile (.9). However, as the predicted “no-go” RT quantiles were still highly correlated with empirical quantiles (*r* >= .72), we concluded that the model described individual differences in “no-go” RTs relatively well despite the low number of “no-go” trials. We also conducted parameter recovery analyses following methods similar to those outlined in recent studies (White et al., 2018), and found that all parameters displayed “good” recovery (*r* between simulated and recovered parameters > .75), except for the z parameter, for which recovery was only “fair” (*r* = .68). Relevant to the current study, we found that both the *v.go* (*r* = .90) and *v.nogo* (*r* = .86) parameters could be effectively recovered from this go/no-go task.

**Correlations Between Raw and Composite Substance Use Measures**

We investigated simple correlations between all individual and composite substance use outcome measures of interest from ages 22–26 in order to 1) evaluate whether they were moderately correlated with each other, as would be expected given prior research, and 2) ensure that our substance use composite (SC) measure was well-representative of the use of all three substances. We also did the same with measures of prior cumulative use of all three substances from age 17 and the prior substance use composite (preSC) that was utilized as a covariate in the primary analyses. Supplemental Table 1 displays Bayesian estimates of correlation coefficients between all of these variables. Inspection of these values indicates that, as expected, measures of average use of all three substances from ages 22–26 are moderately correlated with one another, and measures of cumulative use by age 17 also show strong interrelationships. Furthermore, both the SC and preSC show strong, and roughly equal, correlations with the three substance use measures that each composite measure was derived from, suggesting that these composites provide representative indices of the use of common substances for specific developmental periods.

**False Alarm Rate Analysis**

Although we primarily aimed to test the hypothesis that efficiency of evidence accumulation (EEA) predicts substance use, and therefore focused on the drift rate parameters of the DDM (*v.go*, *v.nogo*, *v.avg*), we also conducted a supplemental prediction analysis using the false alarm (FA) rate, the primary behavioral index of “inhibitory control” from the task. We used the same covariates as in the primary analyses involving DDM parameters. Results from this analysis (Supplemental Table 2) indicate that FA rate was not a statistically significant predictor of the SC. Hence, consistent with results from previous longitudinal studies that aimed to predict substance use with measures drawn from the go/no-go task (Heitzeg et al., 2014; Mahmood et al., 2013; Wetherill et al., 2013), our data suggest that FA rate is not a robust predictor of use.

**Sensitivity Analyses**

To assess whether our findings were robust to the inclusion of prior substance use and other covariates in our prediction models, we conducted two sensitivity analyses using the same frequentist and Bayesian methods that were utilized in the primary analyses. First, we conducted prediction analyses which used measures of cumulative use of individual substances by age 17 (alcohol, marijuana, cigarettes) as covariates, in place of the preSC, in order to evaluate whether our findings would hold when prior use of these substances was accounted for individually (Supplemental Tables 3–5). Next, we conducted prediction analyses without any of our previous covariates to evaluate whether our results were still robust even when models did not account for other relevant risk factors (Supplemental Tables 6 and 7).

Results of the first sensitivity analysis are highly similar to the results of our primary analyses reported in the manuscript; both predictors of interest, *v.avg* and PC1, show statistically significant relationships with the SC outcome in frequentist tests, and Bayesian model comparison indicates substantial evidence for the inclusion of both predictors of interest in the model. Results of our second sensitivity analysis, without any covariates, were also similar, although there was slightly less evidence for the inclusion of the neural-level measure (PC1). However, the best-fitting model was still one which contained both predictors of interest, rather than *v.avg* only. Taken together, results from these sensitivity analyses suggest that our primary results are generally robust to the inclusion, vs. exclusion, of covariates in our regression models, and to alterations in the measurement of the prior substance use covariates, specifically.

**Individual Regions of Interest as Predictors of Substance Use**

We opted to use a network-based approach, in which a latent component score that was informed by activation across multiple regions of interest (ROIs) linked to error monitoring provided our primary measure of error-related activation, for our main analyses due to its advantages relative to traditional univariate approaches. Nonetheless, we also appreciate that readers may be interested in whether activation estimates from individual ROIs display more selective prospective relationships with substance use behaviors. We did not attempt Bayesian model comparison analyses to investigate this possibility because we were concerned that the high degree of collinearity between many pairs of ROIs (*r*>.50) would render models that included these ROIs uninformative. However, to provide preliminary indications of which ROIs may display more selective relationships with substance use, we conducted eight separate frequentist regression analyses involving each individual ROI along with covariates (Supplemental Tables 8–9). These analyses indicated that only the ACC and bilateral insula/IFG ROIs demonstrated prospective relationships with substance use that survived correction for multiple comparisons, although the bilateral parietal ROIs also displayed indications of weaker prospective relationships.

Supplemental Tables

**Supplemental Table 1**. Correlations between measures of the use of individual substances, both averaged over ages 22–26 and cumulative use (Cu.) by age 17, as well as with the age 22–26 average substance use composite (SC) and age 17 prior substance use composite (preSC). Large-font numbers indicate the median of the Bayesian posterior distribution of the correlation coefficient, representing the most likely correlation value, while smaller-font numbers in italics indicate the 95% credible intervals of the posterior distribution, which represent the upper and lower bounds of the range in which there is a .95 probability that the correlation coefficient falls. DV = annual volume of alcoholic drinks (standard beverages); MF = annual marijuana use frequency (days of use); CF = annual cigarette use frequency (days of use)

|  | DV (22–26) | MF (22–26) | CF  (22–26) | SC (22–26) | Cu. DV (17) | Cu. MF (17) | Cu. CF (17) |
| --- | --- | --- | --- | --- | --- | --- | --- |
| DV (22–26) | — |  |  |  |  |  |  |
|  | — |  |  |  |  |  |  |
|  | — |  |  |  |  |  |  |
| MF (22–26) | 0.24 | — |  |  |  |  |  |
|  | *0.41* | — |  |  |  |  |  |
|  | *0.05* | — |  |  |  |  |  |
| CF (22–26) | 0.34 | 0.29 | — |  |  |  |  |
|  | *0.49* | *0.45* | — |  |  |  |  |
|  | *0.16* | *0.10* | — |  |  |  |  |
| SC (22–26) | 0.73 | 0.70 | 0.75 | — |  |  |  |
|  | *0.80* | *0.78* | *0.82* | — |  |  |  |
|  | *0.62* | *0.59* | *0.64* | — |  |  |  |
| Cu. DV (17) | 0.60 | 0.28 | 0.26 | 0.53 | — |  |  |
|  | *0.71* | *0.45* | *0.43* | *0.65* | — |  |  |
|  | *0.46* | *0.10* | *0.08* | *0.37* | — |  |  |
| Cu. MF (17) | 0.40 | 0.39 | 0.24 | 0.47 | 0.71 | — |  |
|  | *0.55* | *0.53* | *0.41* | *0.60* | *0.79* | — |  |
|  | *0.23* | *0.21* | *0.05* | *0.30* | *0.60* | — |  |
| Cu. CF (17) | 0.29 | 0.24 | 0.33 | 0.39 | 0.70 | 0.64 | — |
|  | *0.45* | *0.40* | *0.49* | *0.54* | *0.79* | *0.73* | — |
|  | *0.11* | *0.05* | *0.15* | *0.22* | *0.59* | *0.50* | — |
| preSC (17) | 0.49 | 0.34 | 0.31 | 0.52 | 0.91 | 0.88 | 0.88 |
|  | *0.61* | *0.49* | *0.47* | *0.64* | *0.94* | *0.92* | *0.91* |
|  | *0.32* | *0.16* | *0.13* | *0.36* | *0.86* | *0.83* | *0.82* |

**Supplemental Table 2**. Results from the frequentist regression analysis that attempted to predict values of the age 22–26 substance use composite (SC) with false alarm (FA) rate, the primary behavioral index of inhibition from the go/no-go task, and our standard covariates. **Bolded** *p*-values survive false discovery rate correction for multiple comparisons. Overall variance explained by the model (R^2^) is displayed in parentheses. R/E = Race/Ethnicity; AUD FHx = family history of alcohol use disorder (either parent); ADHD Dx = Attention-Deficit/Hyperactivity Disorder diagnosis; preSC = prior substance use composite (cumulative use through age 17)

| **Model (R^2^)** |  | **Unstandardized** | **Standard Error** | **Standardized** | ***t*** | ***p*** |
| --- | --- | --- | --- | --- | --- | --- |
| **FAs** | (Intercept) | -0.122 | 0.326 |  |  |  |
| **(.355)** | Sex | -0.274 | 0.120 | -0.186 | -2.273 | 0.025 |
|  | R/E | 0.02 | 0.314 | 0.005 | 0.064 | 0.949 |
|  | AUD FHx | 0.017 | 0.135 | 0.01 | 0.123 | 0.902 |
|  | ADHD Dx | 0.314 | 0.243 | 0.108 | 1.294 | 0.199 |
|  | preSC | 0.433 | 0.067 | 0.529 | 6.487 | **< .001** |
|  | FAs | 0.731 | 0.424 | 0.148 | 1.726 | 0.088 |

**Supplemental Table 3**. Results from our first sensitivity analysis, which included measures of prior use of individual substances as covariates rather than a prior use composite, involving frequentist regressions that predicted values of the age 22–26 substance use composite (SC) with models that included 1) *v.go* 2) *v.nogo*, and 3) *v.avg*, along with nuisance covariates. **Bolded** *p*-values survive false discovery rate correction for multiple comparisons within families defined by the individual regression models. Overall variance explained by each model (R^2^) is displayed in parentheses. R/E = Race/Ethnicity; AUD FHx = family history of alcohol use disorder (either parent); ADHD Dx = Attention-Deficit/Hyperactivity Disorder diagnosis; Cu. DV = cumulative drink volume at age 17; Cu. MJ = cumulative marijuana use at age 17; Cu. CF = cumulative cigarette use at age 17

| **Model (R^2^)** |  | **Unstandardized** | **Standard Error** | **Standardized** | ***t*** | ***p*** |
| --- | --- | --- | --- | --- | --- | --- |
| ***v.go*** | (Intercept) | 0.293 | 0.356 |  |  |  |
| **(.404)** | Sex | -0.258 | 0.118 | -0.175 | -2.183 | 0.031 |
|  | R/E | 0.069 | 0.300 | 0.018 | 0.23 | 0.819 |
|  | AUD FHx | 0.077 | 0.134 | 0.047 | 0.575 | 0.567 |
|  | ADHD Dx | 0.337 | 0.235 | 0.116 | 1.434 | 0.155 |
|  | Cu. DV (17) | 3.528e -4 | 1.214e -4 | 0.372 | 2.905 | **0.005** |
|  | Cu. MJ (17) | 0.001 | 6.575e -4 | 0.182 | 1.536 | 0.128 |
|  | Cu. CF (17) | -9.866e -6 | 2.451e -4 | -0.005 | -0.04 | 0.968 |
|  | *v.go* | -0.167 | 0.057 | -0.238 | -2.918 | **0.004** |
| ***v.nogo*** | (Intercept) | 0.132 | 0.362 |  |  |  |
| **(.377)** | Sex | -0.236 | 0.121 | -0.161 | -1.946 | 0.054 |
|  | R/E | 0.046 | 0.309 | 0.012 | 0.148 | 0.883 |
|  | AUD FHx | 0.049 | 0.136 | 0.03 | 0.359 | 0.72 |
|  | ADHD Dx | 0.356 | 0.241 | 0.122 | 1.475 | 0.143 |
|  | Cu. DV (17) | 3.597e -4 | 1.241e -4 | 0.379 | 2.898 | **0.005** |
|  | Cu. MJ (17) | 9.130e -4 | 6.708e -4 | 0.165 | 1.361 | 0.177 |
|  | Cu. CF (17) | 4.506e -5 | 2.492e -4 | 0.022 | 0.181 | 0.857 |
|  | *v.nogo* | -0.15 | 0.075 | -0.166 | -1.986 | 0.050 |
| ***v.avg*** | (Intercept) | 0.286 | 0.363 |  |  |  |
| **(.397)** | Sex | -0.244 | 0.119 | -0.166 | -2.05 | 0.043 |
|  | R/E | 0.044 | 0.303 | 0.012 | 0.145 | 0.885 |
|  | AUD FHx | 0.073 | 0.135 | 0.044 | 0.541 | 0.590 |
|  | ADHD Dx | 0.333 | 0.237 | 0.114 | 1.403 | 0.164 |
|  | Cu. DV (17) | 3.561e -4 | 1.221e -4 | 0.375 | 2.915 | **0.004** |
|  | Cu. MJ (17) | 9.744e -4 | 6.608e -4 | 0.176 | 1.474 | 0.144 |
|  | Cu. CF (17) | 7.822e -6 | 2.460e -4 | 0.004 | 0.032 | 0.975 |
|  | *v.avg* | -0.187 | 0.069 | -0.222 | -2.695 | **0.008** |

**Supplemental Table 4**. Results from our first sensitivity analysis, which included measures of prior use of individual substances as covariates rather than a prior use composite, involving frequentist regressions that predicted values of the age 22–26 substance use composite (SC) with models that included 1) PC1 and 2) PC2, along with nuisance covariates. **Bolded** *p*-values survive false discovery rate correction for multiple comparisons within families defined by the individual regression models. Overall variance explained by each model (R^2^) is displayed in parentheses. R/E = Race/Ethnicity; AUD FHx = family history of alcohol use disorder (either parent); ADHD Dx = Attention-Deficit/Hyperactivity Disorder diagnosis; Cu. DV = cumulative drink volume at age 17; Cu. MJ = cumulative marijuana use at age 17; Cu. CF = cumulative cigarette use at age 17

| **Model (R^2^)** |  | **Unstandardized** | **Standard Error** | **Standardized** | ***t*** | ***p*** |
| --- | --- | --- | --- | --- | --- | --- |
| **PC1** | (Intercept) | -0.323 | 0.308 |  |  |  |
| **(.418)** | Sex | -0.205 | 0.118 | -0.139 | -1.734 | 0.086 |
|  | R/E | 0.175 | 0.296 | 0.046 | 0.59 | 0.556 |
|  | AUD FHx | 0.089 | 0.132 | 0.054 | 0.673 | 0.502 |
|  | ADHD Dx | 0.323 | 0.233 | 0.111 | 1.388 | 0.168 |
|  | Cu. DV (17) | 3.905e -4 | 1.204e -4 | 0.411 | 3.243 | **0.002** |
|  | Cu. MJ (17) | 9.672e -4 | 6.490e -4 | 0.174 | 1.490 | 0.139 |
|  | Cu. CF (17) | 1.547e -5 | 2.411e -4 | 0.007 | 0.064 | 0.949 |
|  | PC1 | -0.092 | 0.028 | -0.268 | -3.314 | **0.001** |
| **PC2** | (Intercept) | -0.22 | 0.319 |  |  |  |
| **(.370)** | Sex | -0.297 | 0.124 | -0.202 | -2.408 | 0.018 |
|  | R/E | 0.185 | 0.31 | 0.049 | 0.598 | 0.551 |
|  | AUD FHx | -0.032 | 0.137 | -0.02 | -0.235 | 0.815 |
|  | ADHD Dx | 0.367 | 0.242 | 0.126 | 1.515 | 0.133 |
|  | Cu. DV (17) | 3.851e -4 | 1.259e -4 | 0.406 | 3.059 | **0.003** |
|  | Cu. MJ (17) | 0.001 | 6.812e -4 | 0.187 | 1.52 | 0.132 |
|  | Cu. CF (17) | -3.385e -5 | 2.590e -4 | -0.016 | -0.131 | 0.896 |
|  | PC2 | 0.081 | 0.049 | 0.147 | 1.674 | 0.097 |

**Supplemental Table 5.** Results from our first sensitivity analysis, which included measures of prior use of individual substances as covariates rather than a prior use composite, where Bayesian regression models that involved all combinations of predictors of interest, drift rate (*v.avg*) and error-related activation (PC1), were compared to a “null” model that included only the covariates of sex, race/ethnicity (R/E) family history of alcohol use disorder (AUD FHx), Attention-Deficit/Hyperactivity Disorder diagnosis (ADHD), and separate measures of prior alcohol (DV), marijuana (MJ) and cigarette use (CF). In the “Model Comparison” section, P(M) is prior probability of the model, P(M|data) is the posterior probability of the model after seeing the data, BF_10_ is the Bayes factor comparing the model to the “null” model, and BF_M_ is a Bayes factor comparing the model to all other models from the analysis. The “Posterior Summaries” section reports the model-averaged mean, standard deviation (SD) and 95% credible intervals of posterior samples for coefficients of each predictor of interest, as well as inclusion probabilities obtained from model averaging; P(inc) is the prior probability of including each predictor, P(inc|data) is the posterior probability of including each predictor, and BF_inc_ is a Bayes factor for the change from prior to posterior inclusion odds for the predictor after seeing the data.

| **Model Comparison** | | | | | | | | | | | |
| --- | --- | --- | --- | --- | --- | --- | --- | --- | --- | --- | --- |
| **Models** | | **P(M)** | | **P(M\|data)** | | | **BF_M_** | | **BF_10_** | | ***R*²** |
| “Null” (Sex, R/E, AUD FHx, ADHD, DV, MJ, CF) |  | 0.125 |  | | 0.008 |  | 0.025 |  | 1 |  | 0.352 |
| *v.avg* + PC1 |  | 0.125 |  | | 0.650 |  | 5.576 |  | 78.860 |  | 0.444 |
| PC1 |  | 0.125 |  | | 0.280 |  | 1.169 |  | 33.999 |  | 0.418 |
| *v.avg* |  | 0.125 |  | | 0.061 |  | 0.196 |  | 7.425 |  | 0.397 |

| **Posterior Summaries of Coefficients** | | | | | | | | | | | | | | | |
| --- | --- | --- | --- | --- | --- | --- | --- | --- | --- | --- | --- | --- | --- | --- | --- |
|  | | | | | | | | | | | | | **95% Credible Interval** | | |
| **Coefficient** | | **Mean** | | | **SD** | | **P(inc)** | | **P(inc\|data)** | | **BF_inc_** | | **Lower** | | **Upper** |
| *v.avg* | |  | -0.095 |  | 0.082 |  | 0.500 |  | 0.711 |  | 2.465 |  | -0.241 |  | 0.000 |
| PC1 | |  | -0.069 |  | 0.032 |  | 0.500 |  | 0.931 |  | 13.396 |  | -0.116 |  | 1.400e -5 |

**Supplemental Table 6**. Results from our second sensitivity analysis, which included no covariates for other substance use risk factors, involving frequentist regressions that attempted to predict values of the age 22–26 substance use composite (SC) with models that included 1) *v.go*, 2) *v.nogo*, 3) *v.avg*, 4) PC1, and 5) PC2. **Bolded** *p*-values survive false discovery rate correction for multiple comparisons within families defined by the individual regression models. Overall variance explained by each model (R^2^) is displayed in parentheses.

| **Model (R^2^)** |  | **Unstandardized** | **Standard Error** | **Standardized** | ***t*** | ***p*** |
| --- | --- | --- | --- | --- | --- | --- |
| ***v.go*** | (Intercept) | 0.594 | 0.206 |  |  |  |
| **(.082)** | *v.go* | -0.201 | 0.066 | -0.286 | -3.047 | **0.003** |
| ***v.nogo*** | (Intercept) | 0.410 | 0.189 |  |  |  |
| **(.050)** | *v.nogo* | -0.201 | 0.086 | -0.223 | -2.330 | **0.022** |
| ***v.avg*** | (Intercept) | 0.582 | 0.210 |  |  |  |
| **(.076)** | *v.avg* | -0.233 | 0.080 | -0.276 | -2.931 | **0.004** |
| **PC1** | (Intercept) | 0.020 | 0.069 |  |  |  |
| **(.059)** | PC1 | -0.083 | 0.033 | -0.243 | -2.554 | **0.012** |
| **PC2** | (Intercept) | 0.002 | 0.071 |  |  |  |
| **(.004)** | PC2 | 0.034 | 0.054 | 0.061 | 0.620 | 0.536 |

**Supplemental Table 7**. Results from our second sensitivity analysis, which included no covariates for other substance use risk factors, involving Bayesian regression analyses in which all possible models involving predictors of interest, drift rate (*v.avg*) and error-related activation (PC1), were compared to a “null” model that included only the regression intercept parameter. In the “Model Comparison” section, P(M) is prior probability of the model, P(M|data) is the posterior probability of the model after seeing the data, BF_10_ is the Bayes factor comparing the model to the “null” model, and BF_M_ is a Bayes factor comparing the model to all other models from the analysis. The “Posterior Summaries” section reports the model-averaged mean, standard deviation (SD) and 95% credible intervals of posterior samples for coefficients of each predictor of interest, as well as inclusion probabilities obtained from model averaging; P(inc) is the prior probability of including each predictor, P(inc|data) is the posterior probability of including each predictor, and BF_inc_ is a Bayes factor for the change from prior to posterior inclusion odds for the predictor after seeing the data.

| **Model Comparison** | | | | | | | | | | | |
| --- | --- | --- | --- | --- | --- | --- | --- | --- | --- | --- | --- |
| **Models** | | **P(M)** | | **P(M\|data)** | | | **BF_M_** | | **BF_10_** | | **R²** |
| “Null” (intercept only) |  | 0.250 |  | | 0.040 |  | 0.124 |  | 1.000 |  | 0.000 |
| *v.avg* + PC1 |  | 0.250 |  | | 0.472 |  | 2.680 |  | 11.903 |  | 0.109 |
| *v.avg* |  | 0.250 |  | | 0.346 |  | 1.584 |  | 8.717 |  | 0.076 |
| PC1 |  | 0.250 |  | | 0.143 |  | 0.500 |  | 3.605 |  | 0.059 |

| **Posterior Summaries of Coefficients** | | | | | | | | | | | | | | |
| --- | --- | --- | --- | --- | --- | --- | --- | --- | --- | --- | --- | --- | --- | --- |
|  | | | | | | | | | | | | **95% Credible Interval** | | |
| **Coefficient** | **Mean** | | | **SD** | | **P(inc)** | | **P(inc\|data)** | | **BF_inc_** | | **Lower** | | **Upper** |
| *v.avg* |  | -0.156 |  | 0.103 |  | 0.500 |  | 0.817 |  | 4.478 |  | -0.317 |  | 0.000 |
| PC1 |  | -0.038 |  | 0.039 |  | 0.500 |  | 0.615 |  | 1.596 |  | -0.114 |  | 0.000 |

**Supplemental Table 8**. Results from frequentist regression analyses predicting values of the age 22–26 substance use composite (SC) with models that included activation estimates from 1) anterior cingulate cortex (ACC), 2) left insula / inferior frontal gyrus (LI/IFG), 3) right insula / inferior frontal gyrus (RI/IFG), and 4) the pre-supplementary motor area (PSMA), along with covariates. **Bolded** *p*-values survive false discovery rate correction for multiple comparisons within families defined by the individual regression models. Overall variance explained by each model (R^2^) is displayed in parentheses. R/E = Race/Ethnicity; AUD FHx = family history of alcohol use disorder (either parent); ADHD Dx = Attention-Deficit/Hyperactivity Disorder diagnosis; preSC = prior substance use composite (cumulative use through age 17)

| **Model (R^2^)** |  | **Unstandardized** | **Standard Error** | **Standardized** | ***T*** | ***p*** |
| --- | --- | --- | --- | --- | --- | --- |
| **ACC** | (Intercept) | 0.143 | 0.322 |  |  |  |
| **(.375)** | Sex | -0.243 | 0.12 | -0.165 | -2.033 | 0.045 |
|  | R/E | 0.076 | 0.304 | 0.02 | 0.249 | 0.804 |
|  | AUD FHx | 0.021 | 0.132 | 0.013 | 0.162 | 0.872 |
|  | ADHD Dx | 0.369 | 0.234 | 0.127 | 1.572 | 0.119 |
|  | preSC | 0.435 | 0.066 | 0.532 | 6.631 | **< .001** |
|  | ACC | -0.054 | 0.022 | -0.202 | -2.47 | **0.015** |
| **LI/IFG** | (Intercept) | 0.117 | 0.304 |  |  |  |
| **(.424)** | Sex | -0.329 | 0.114 | -0.223 | -2.898 | **0.005** |
|  | R/E | 0.18 | 0.292 | 0.047 | 0.617 | 0.539 |
|  | AUD FHx | 0.063 | 0.128 | 0.039 | 0.495 | 0.621 |
|  | ADHD Dx | 0.208 | 0.230 | 0.071 | 0.905 | 0.368 |
|  | preSC | 0.434 | 0.063 | 0.531 | 6.907 | **< .001** |
|  | LI/IFG | -0.089 | 0.023 | -0.310 | -3.889 | **< .001** |
| **RI/IFG** | (Intercept) | 0.14 | 0.311 |  |  |  |
| **(.404)** | Sex | -0.323 | 0.115 | -0.22 | -2.8 | **0.006** |
|  | R/E | 0.204 | 0.297 | 0.054 | 0.686 | 0.494 |
|  | AUD FHx | 0.02 | 0.129 | 0.012 | 0.157 | 0.876 |
|  | ADHD Dx | 0.242 | 0.233 | 0.083 | 1.04 | 0.301 |
|  | preSC | 0.428 | 0.064 | 0.523 | 6.697 | **< .001** |
|  | RI/IFG | -0.079 | 0.024 | -0.269 | -3.367 | **0.001** |
| **PSMA** | (Intercept) | -0.007 | 0.324 |  |  |  |
| **(.341)** | Sex | -0.291 | 0.121 | -0.198 | -2.405 | 0.018 |
|  | R/E | 0.142 | 0.312 | 0.037 | 0.454 | 0.651 |
|  | AUD FHx | 0.007 | 0.138 | 0.004 | 0.051 | 0.959 |
|  | ADHD Dx | 0.393 | 0.240 | 0.135 | 1.635 | 0.105 |
|  | preSC | 0.432 | 0.068 | 0.529 | 6.339 | **< .001** |
|  | PSMA | -0.02 | 0.024 | -0.073 | -0.855 | 0.395 |

**Supplemental Table 9**. Results from frequentist regression analyses predicting values of the age 22–26 substance use composite (SC) with models that included activation estimates from 1) left parietal lobe (LPar), 2) right parietal lobe (RPar), 3) left striatum (LStri), and 4) right striatum (RStri), along with covariates. **Bolded** *p*-values survive false discovery rate correction for multiple comparisons within families defined by the individual regression models. Overall variance explained by each model (R^2^) is displayed in parentheses. R/E = Race/Ethnicity; AUD FHx = family history of alcohol use disorder (either parent); ADHD Dx. = Attention-Deficit/Hyperactivity Disorder diagnosis; preSC = prior substance use composite (cumulative use through age 17)

| **Model (R^2^)** |  | **Unstandardized** | **Standard Error** | **Standardized** | ***T*** | ***p*** |
| --- | --- | --- | --- | --- | --- | --- |
| **LPar** | (Intercept) | 0.01 | 0.316 |  |  |  |
| **(.373)** | Sex | -0.241 | 0.12 | -0.164 | -2.008 | 0.047 |
|  | R/E | 0.218 | 0.306 | 0.057 | 0.711 | 0.479 |
|  | AUD FHx | -0.006 | 0.132 | -0.003 | -0.043 | 0.965 |
|  | ADHD Dx | 0.338 | 0.236 | 0.116 | 1.434 | 0.155 |
|  | preSC | 0.43 | 0.066 | 0.526 | 6.556 | **< .001** |
|  | LPar | -0.079 | 0.033 | -0.199 | -2.421 | 0.017 |
| **RPar** | (Intercept) | 0.045 | 0.317 |  |  |  |
| **(.370)** | Sex | -0.26 | 0.119 | -0.177 | -2.178 | 0.032 |
|  | R/E | 0.188 | 0.306 | 0.050 | 0.615 | 0.540 |
|  | AUD FHx | -0.016 | 0.132 | -0.010 | -0.119 | 0.905 |
|  | ADHD Dx | 0.357 | 0.236 | 0.123 | 1.513 | 0.133 |
|  | preSC | 0.424 | 0.066 | 0.519 | 6.464 | **< .001** |
|  | RPar | -0.077 | 0.033 | -0.188 | -2.314 | 0.023 |
| **LStri** | (Intercept) | -0.035 | 0.326 |  |  |  |
| **(.339)** | Sex | -0.272 | 0.127 | -0.185 | -2.151 | 0.034 |
|  | R/E | 0.104 | 0.314 | 0.027 | 0.331 | 0.742 |
|  | AUD FHx | 0.004 | 0.14 | 0.003 | 0.030 | 0.976 |
|  | ADHD Dx | 0.409 | 0.241 | 0.140 | 1.695 | 0.093 |
|  | preSC | 0.424 | 0.067 | 0.519 | 6.297 | **< .001** |
|  | LStri | -0.033 | 0.054 | -0.054 | -0.608 | 0.545 |
| **RStri** | (Intercept) | -0.053 | 0.329 |  |  |  |
| **(.339)** | Sex | -0.278 | 0.124 | -0.189 | -2.252 | 0.027 |
|  | R/E | 0.129 | 0.312 | 0.034 | 0.413 | 0.680 |
|  | AUD FHx | 5.555e -4 | 0.138 | 3.389e -4 | 0.004 | 0.997 |
|  | ADHD Dx | 0.4 | 0.241 | 0.138 | 1.663 | 0.099 |
|  | preSC | 0.42 | 0.067 | 0.514 | 6.233 | **< .001** |
|  | RStri | -0.035 | 0.054 | -0.055 | -0.650 | 0.517 |
